# Comprehensive Analysis Reveals Adaptive DNA Repair and Replication Stress Networks in Genomically Unstable Breast Cancer

**DOI:** 10.64898/2026.05.13.724208

**Authors:** Farah Ramadan, Sara Zahraeifard, Neelima Yadav, Ujwal Subedi, Sameer Shah, Arvind Panday

## Abstract

Genomic instability is a defining hallmark of breast cancer, yet the mechanisms by which tumors tolerate persistent DNA damage remain poorly understood. We performed a comprehensive, multi-cohort analysis of breast cancer datasets to define how DNA damage response (DDR) and replication stress tolerance (RST) networks are rewired in genomically unstable tumors. Using fraction of genome altered (FGA) as a chromosomal instability metric, we show that BRCA-mutant tumors exhibit elevated genomic instability coupled with increased expression of homologous recombination, Fanconi anemia, mismatch repair, base excision repair, and alternative end-joining pathways. Strikingly, heightened pathway activity correlates with increased genome alteration, supporting a model of damage tolerance rather than repair restoration. RST programs, including fork remodeling, protection, and single strand DNA gap suppression, further contribute to tumor fitness under replication stress. These adaptive states are enriched in aggressive subtypes, intensified with progression, and associate with pathway-specific mutational burden. Co-occurrence and mutual exclusivity mapping uncovered non-random subtype-relevant genetic interactions states among major drivers and DDR genes, nominating context-specific synthetic lethal opportunities. Our findings identify compensatory genome-maintenance programs as central drivers of tumor resilience and highlight pathway-specific vulnerabilities for targeted therapeutic intervention.

## INTRODUCTION

Breast cancer (BC) is a biologically heterogeneous disease affecting women globally and characterized by diverse genomic alterations, molecular markers, and gene-expression profiles^1,2^. Molecular classification informs prognostication and therapeutic decision-making; tumors expressing estrogen and/or progesterone receptors (ER/PR) are generally responsive to endocrine therapies, while overexpression of the ERBB2 gene (HER2) defines a distinct subtype that benefits from HER2-targeted treatments^3,4^. Breast cancers lacking ER, PR, and HER2 expression—known as triple-negative breast cancer (TNBC)—are more aggressive, display poorer outcomes and are not eligible for these targeted therapies^5,6^. Based on gene-expression and molecular characteristics, BC is commonly categorized into four intrinsic subtypes^1^. Luminal A tumors are ER-positive, PR-positive, HER2-negative, typically showing slower growth and the most favorable prognosis. Luminal B tumors are also ER-positive but often display lower PR expression, higher proliferation levels, and occasional HER2 positivity, resulting in a more aggressive phenotype that may require combined endocrine therapy, chemotherapy, and anti-HER2 treatment. The HER2-enriched subtype is characterized by high HER2 expression and low ER/PR levels, tends to be highly proliferative, but responds well to HER2-directed therapies. The basal-like subtype substantially overlaps with TNBC and is typically composed of high-grade, aggressive tumors that occur more frequently in younger women and individuals with *BRCA1* mutations, with chemotherapy remaining the main treatment option.

Genomic instability is a defining hallmark of BC and is widely considered a critical driver for genetic alterations and subsequent malignant transformation^7,8^. This instability often arises early during carcinogenesis as a consequence of defects in genome “caretaker” functions, including pathways responsible for cell cycle control, replication stress tolerance (RST), DNA damage detection, and DNA repair^9^. This exposes actionable liabilities, particularly in DNA repair and RST pathways, that can be therapeutically exploited. Yet, despite widespread repair and RST defects, established tumors frequently maintain robust tolerance to genotoxic stress. Converging evidence indicates that this resilience is mediated by dynamic cross-talk among multiple DNA repair and RST pathways, whereby loss of one mechanism is buffered by compensatory activation of others^10^.

BRCA1 and BRCA2 are essential tumor suppressor proteins that function as core regulators of genome stability by orchestrating homologous recombination repair (HRR), a high-fidelity DNA repair pathway responsible for the repair of DNA double-strand breaks (DSBs). BRCA1 contributes to the early stages of repair by promoting DNA end resection and coordinating the recruitment of downstream repair factors, while BRCA2 plays a central role in the later stages by facilitating the loading of RAD51 onto single-stranded DNA to drive homology search and strand invasion^11,12^. Defects in HRR, referred as homologous recombination deficiency (HRD), represent a major source of genomic instability and a key driver of tumorigenesis^13,14^. In BC, HRD is associated with therapeutic vulnerability to poly(ADP-ribose) polymerase inhibitors (PARPi), which exploit synthetic lethality in HR-deficient cells^15,16^. Beyond germline or somatic mutations in *BRCA1* and *BRCA2*, the concept of “BRCAness” has been introduced to describe tumors that phenocopy key molecular and functional defects of *BRCA1/2-*mutant cancers despite retaining intact *BRCA* genes^17^. A defining feature of BRCAness is heightened sensitivity to PARP inhibition, traditionally attributed to impaired HR-mediated DNA repair.

Beyond their canonical role in HRR, BRCA1 and BRCA2 are also key mediators of RST mechanisms. A central RST mechanism is replication fork reversal. BRCA1 facilitates this process through coordination with fork remodeling enzymes, such as RAD51 and SMARCAL1, and, together with BRCA2, protects reversed fork arms from nucleolytic degradation^18–21^. When fork reversal is impaired, *BRCA*-mutant cells rely on PRIMPOL-dependent repriming, which promotes accumulation of single-stranded DNA (ssDNA) gaps; notably, ssDNA gap burden correlates with chemotherapy response in *BRCA*-mutant tumors, underscoring the clinical relevance of this pathway^22^. BRCA1 also suppresses group-1 tandem duplications (TDs), a characteristic repair outcome that arises in response to replication stress. These TDs are defined by head-to-tail duplications of chromosomal segments within the same rearranged chromosome and represent a distinctive class of structural variants that are frequently observed in BC^23^. Therefore, loss of BRCA1 or BRCA2 function not only leads to profound defects in HRR but also accumulates replication stress which contributes to chromosome rearrangement and genomic instability. Despite these vulnerabilities, *BRCA*-mutant breast tumors frequently retain the capacity to survive and proliferate under high levels of genotoxic stress. This paradox suggests that *BRCA*-mutant tumors undergo adaptive rewiring of the DNA damage response (DDR) and RST pathways, engaging compensatory repair mechanisms that sustain genome integrity and tumor viability. Therefore, systematically mapping compensatory repair circuits across breast tumor grades and molecular subtypes is critical to transform BRCA-associated genomic instability from a tolerated state into a clinically actionable lethal vulnerability.

Tumorigenesis is a multistep evolutionary process that frequently arises through the co-occurrence of alterations in multiple DNA repair pathways, in which distinct defects accumulate over time and functionally interact to compromise complementary layers of genome maintenance^13,24–28^. From a therapeutic perspective, the presence of co-occurring driver mutations has profound implications for treatment response^29,30^. Analyzing co-occurrence patterns among DNA repair and oncogenic mutations provides valuable insights into tumor risk stratification and resistance prediction. While patterns of co-occurring mutations highlight how complementary defects collaborate to promote tumor development and therapeutic resistance, an equally informative genomic signature arises from the opposite pattern, mutual exclusivity, which often reflects functional redundancy between essential genome maintenance pathways, such that the simultaneous loss of both functions is catastrophic for cancer cell viability^31,32^. This provides a powerful framework for identifying synthetic lethal gene pairs and has guided the development of targeted cancer therapies. A landmark clinical success of this strategy is exemplified by the PARPi. However, their clinically efficacy remains limited, with over 40% of *BRCA*-mutant patients failing to respond to therapy^33,34^. Systematically investigating the mutual exclusivity of DNA repair genes provides a compelling and rational strategy to uncover novel synthetic lethal interactions and therapeutic targets that can overcome PARPi resistance.

Alterations in DDR pathways can also shape tumor immunogenicity. A key biomarker in this context is tumor mutational burden (TMB), defined as the number of somatic nonsynonymous mutations within a defined genomic region, which has been associated with improved response to immune checkpoint inhibitors (ICIs) across cancers, especially TNBC^35,36^. Mechanistically, DDR defects can increase mutation load and neoantigen generation, while also activating innate immune pathways such as cGAS–STING, thereby promoting interferon signaling, antigen presentation, and immune-cell infiltration^37,38^. Consistent with this biology, preclinical studies show that inhibition of DDR targets can enhance tumor immunogenicity and sensitize tumors to ICIs^39–41^. However, the relationship between specific DDR alterations and TMB-linked immunotherapy response in BC remains incompletely defined, in part because DDR alterations are heterogeneous across subtypes and grades, and not all defects produce equivalent immunogenic effects. Defining pathway-specific DDR alterations and their association with TMB is therefore essential for identifying DDR-driven immune phenotypes, improving patient stratification, and guiding rational DDR–ICI combination strategies in BC.

In this study, we perform a comprehensive analysis using publicly available BC patient data from cBioPortal for Cancer genomics (cbioportal.org)^42–44^, an open-access web-based platform to explore and analyze large-scale cancer genomics datasets. We focused on DDR and RST gene expression using The Cancer Genome Atlas (TCGA) invasive breast carcinoma datasets. We systematically define how *BRCA*-associated genomic instability is accommodated by adaptive rewiring of DDR and RST networks in BC. By investigating fraction genome altered as a genome-wide instability metric across independent cohorts in the context of *BRCA1/2*-mutation, we identify important DDR and RST pathways that are activated as compensatory mechanisms that potentially maintain BC cell viability in the face of increased genome instability, providing better understanding of treatment resistance and highlighting new vulnerabilities that can be exploited for treatment. We further investigate how these adaptive tumors associate with tumor progression including advanced stage and aggressive subtypes, linking repair-state plasticity to clinically aggressive disease. Focusing on mutation counts to reveal pathway-specific mutational consequences, we investigate how mutations in different DDR and RST genes contribute to the overall TMB in BC patients. Additionally, co-occurrence and mutual-exclusivity mapping uncovers non-random, gene-specific interaction states among major drivers and DDR pathways, highlighting heterogeneous repair dependencies. Together, these findings establish a framework in which breast tumors exploit distinct compensatory genome-maintenance circuits to tolerate persistent genomic stress, nominating subtype- and pathway-resolved vulnerabilities for rational patient stratification and combination therapeutic design.

## RESULTS

### Genome instability is associated with compensatory DNA repair pathway adaptation in *BRCA*-mutant breast tumors

One widely used metric for characterizing genomic alterations associated with cancer initiation and progression is the Fraction of Genome Altered (FGA) which represents the proportion of the genome affected by somatic copy number variations (CNVs), encompassing both copy number gains and losses of DNA segments. As a quantitative indicator of genomic instability, higher FGA values reflect a greater burden of chromosomal aberrations often associated with disease progression, tumor aggressiveness, treatment resistance, and poorer clinical outcomes^45,46^. To evaluate the effect of *BRCA1* and *BRCA2* loss on genome-wide genomic instability, we analyzed their association with FGA across independent Breast Invasive Carcinoma cohorts: the “MSK 2025”, “TCGA *PanCancer Atlas*”, and the “TCGA *Nature 2012*”^47^ studies publicly available on cbioportal.org. Our analysis revealed that breast tumors harboring *BRCA1* mutation consistently exhibited significantly higher FGA values compared with tumors without *BRCA1* mutation, indicating increased genomic instability **(Fig.1A)**. This association was observed uniformly across all three cohorts, underscoring the reliability of the finding. A similar trend was also evident for *BRCA2* where loss of gene function was likewise associated with elevated FGA levels, but results did not reach significance likely due to low number of *BRCA2* mutant samples **(Fig. S1A).** These results highlight the critical role of *BRCA1* and *BRCA2* in maintaining genomic integrity.

**Figure 1.**
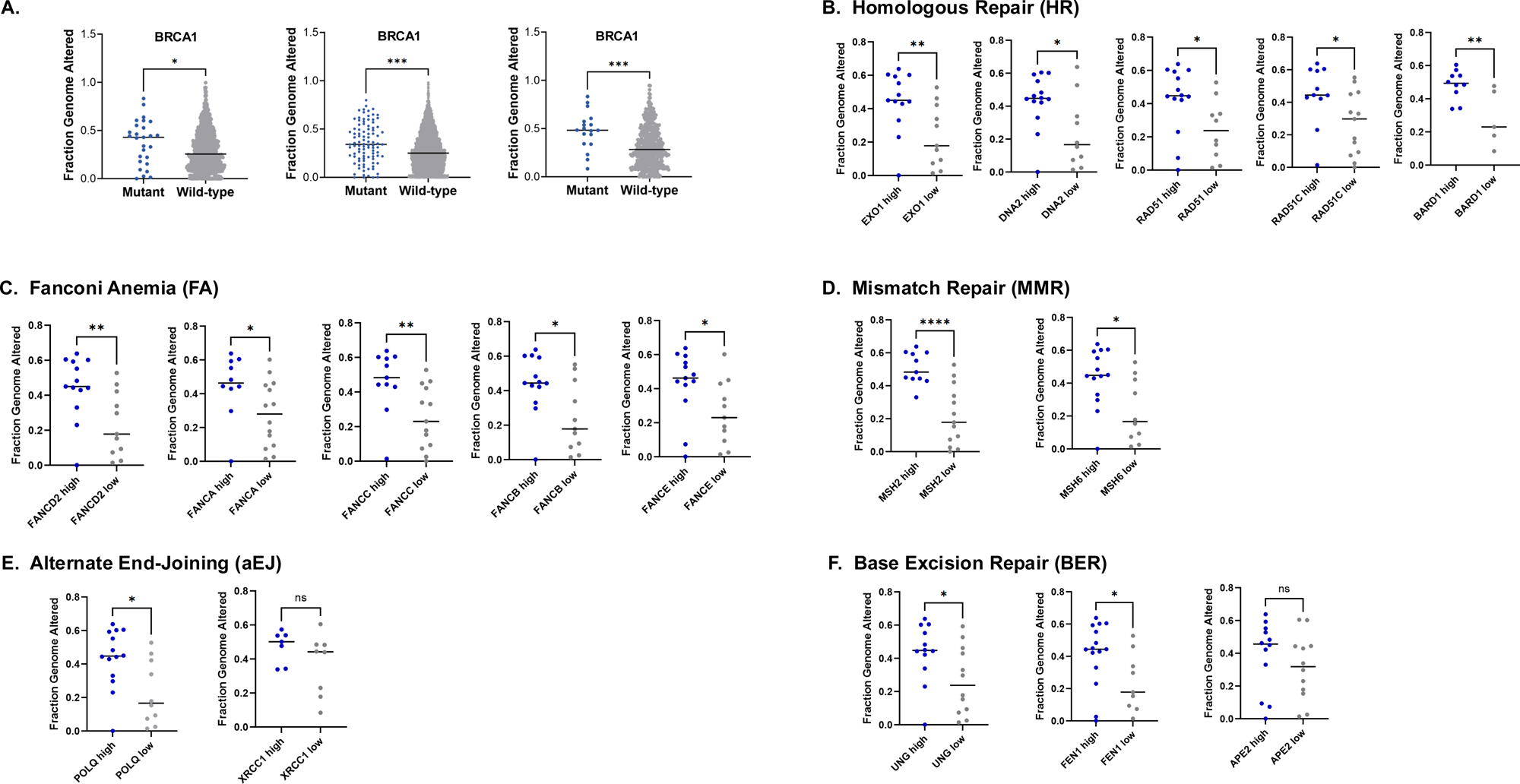
BRCA1 deficiency drives genomic instability and adaptive DNA repair pathway activation in breast tumors. **(A)** FGA levels in TCGA *PanCancer Atlas* (n = 1050) (left), *MSK 2025* (n = 3841) (middle), TCGA *Nature 2012* (n = 482) (right) cohorts relative to *BRCA1* gene status (Wild-type/Mutant). **(B-F)** Expression of DNA-repair genes (high/low) relative to FGA in *BRCA1*-mutant breast cancer samples using TCGA *PanCancer Atlas* (n = 24) or TCGA *Nature 2012* (n = 15) (for BARD1 and XRCC1 genes); genes are grouped into different DNA-repair pathways including **(B)** Homologous repair (HR), **(C)** Fanconi Anemia (FA), **(D)** Mismatch repair, **(E)** Alternative end-joining (aEJ) or **(F)** Base Excision repair (BER). Heatmap mRNA expression z-score <0 was considered as low expression and heatmap mRNA expression z-score >0 was considered as high expression. Asterisks denote significant p-value using two-tailed unpaired t-test with ***p<0.001, **p<0.01, *p<0.05; ns = non-significant.

To explore the adaptive rewiring of the DDR pathways as compensatory repair mechanisms in *BRCA*-mutant breast tumors that sustain genome integrity at levels compatible with tumor viability, we first sought to identify genes within different HRR-associated pathways whose expression levels were associated with FGA. BRCA1 promotes DNA end resection and facilitates homology search by forming a complex with PALB2, which recruits BRCA2 to load RAD51 onto single-stranded DNA^12,48^. Stratification by long-range resection factors revealed a significant increase in FGA in *BRCA1*-mutant tumors with high *EXO1* or *DNA2* expression compared with low-expression counterparts **(Fig.1B)**. A similar pattern was observed for genes mediating homology search, as *RAD51*-high and *RAD51C*-high tumors displayed significantly elevated FGA relative to low-expression groups **(Fig.1B)**. Together, these results indicate that increased expression of both resection– and homology search–associated HR genes in *BRCA1*-mutant tumors correlates with moderate to high levels of chromosomal instability.

We extended this investigation in *BRCA2*-mutant breast tumors and observed a similar trend **(Fig. S1B).** In addition, *BARD1*(critical partner BRCA1)-high tumors exhibited a significant increase in FGA compared with *BARD1*-low tumors, suggesting a potential BRCA1-independent role for BARD1 in promoting tolerance to genomic stress **(Fig.1B)**. We further examined whether non-homologous end joining (NHEJ) pathway components were associated with genomic alteration as an alternative repair mechanism in *BRCA1* or *BRCA2*-mutant tumors, but data revealed no significant differences in FGA across any NHEJ components analyzed **(Fig. S1C)**. Collectively, these findings suggest that upregulation of specific HR-associated factors in *BRCA*-mutant breast tumors is linked to elevated FGA levels, indicating activation of adaptive repair mechanisms.

Fanconi anemia (FA) is a chromosomal instability syndrome caused by repair defects of DNA interstrand crosslinks (ICLs). Recent studies have highlighted extensive crosstalk between FA and HR pathways, with key HR factors such as RAD51 and BRCA1 playing essential roles in ICL repair^49,50^. We therefore investigated FA pathway genes that preserve tumor fitness in *BRCA*-mutant context. Focusing on FANCD2, a central effector of the FA pathway^51^, in *BRCA1*-mutant samples, we found a significant difference in genomic instability between *FANCD2*-high groups, which clustered at moderate to high FGA values, and *FANCD2*-low expressing groups which exhibited significantly reduced FGA levels **(Fig.1C)**. Consistently, high expression of each additional FA core complex gene (FANCA, FANCC, FANCB, and FANCE) was significantly associated with increased FGA **(Fig.1C)**. On the other hand, although high *FANCD2* expression did not significantly correlate with FGA in *BRCA2-*mutant tumors, elevated expression of other FA core complex components remained significantly associated with increased FGA, similar to the pattern observed in *BRCA1-*mutant tumors **(Fig. S1D).** These findings indicate that upregulation of FA pathway genes in *BRCA*-mutant breast tumors does not correspond to restoration of genome stability and is instead associated with heightened chromosomal instability, suggesting a tolerance of ongoing genomic stress rather than preventing its accumulation.

We next investigated the DNA mismatch repair (MMR) associated gene network dependencies in maintaining the fitness of *BRCA1*-mutant breast tumors by buffering otherwise lethal genomic instability. Our analysis focused on the core MMR genes *MSH2* and *MSH6*, which together form the MutSα complex responsible for recognizing and initiating repair at base–base mismatches and small insertion/deletion loops^52^. Tumors with high expression of either gene clustered predominantly within moderate-to-high FGA ranges, while tumors with low expression of both genes exhibited significantly reduced FGA **(Fig.1D)**. We further extended this analysis to *BRCA2*-mutant breast tumors where *MSH2* expression displayed a similar positive association with increased FGA but *MSH6* expression did not reach statistical significance **(Fig. S1E)**. Together, these results highlight pathway-specific yet gene-selective differences in MMR engagement between *BRCA1*- and *BRCA2*-deficient tumors.

We next examined alternative end-joining (aEJ), focusing on two key components: POLQ (DNA polymerase theta), which mediates microhomology-dependent gap filling, and XRCC1, a scaffold protein that coordinates repair through DNA ligase III^53–55^. *BRCA1-* and *BRCA2-*mutant tumors with high expression of *POLQ* exhibited moderate-to-high FGA, whereas tumors with *POLQ*-low expression showed markedly reduced FGA **(Fig. 1E and Fig. S1F)**; however, there was no significant association between high FGA and high *XRCC1* for either *BRCA1* or *BRCA2*-mutant tumors **(Fig. 1E and S1F)**

Given the established role of PARP proteins in base excision repair (BER) and the clinical efficacy of PARP inhibitors, BER also represents a potential therapeutic vulnerability in BRCA-deficient contexts^56,57^. We therefore evaluated BER-associated genes that may support genome maintenance and preserve tumor fitness in this context. Our analysis focused on three core BER components: UNG, FEN1, and APE2^58^. UNG excises uracil residues arising from dUTP misincorporation or cytosine deamination, thereby resolving U:G mismatches. FEN1 is a structure-specific endonuclease that cleaves 5′ DNA flap intermediates during BER, while APE2 functions as a 3′–5′ exonuclease and 3′-phosphodiesterase involved in processing DNA flaps and damaged DNA termini^59^.

*BRCA1*- and *BRCA2*-mutant tumors were stratified by their expression levels of UNG, FEN1, and APE2 and FGA was assessed. In *BRCA1*-mutant tumors, high expression of *UNG* and *FEN1* was significantly associated with moderate-to-high FGA values **(Fig. 1F)**. In contrast, *APE2* expression in *BRCA1-*mutant tumors did not show a statistically significant association with FGA. In *BRCA2*-mutant tumors, elevated expressions of *FEN1* and *APE2* correlated within higher FGA **(Fig. S1G)**, whereas UNG expression did not reach statistical significance. Together, these findings reveal gene-specific engagement of the BER pathway that differs between *BRCA1*- and *BRCA2*-deficient tumors, underscoring distinct compensatory repair dependencies.

Collectively, these findings suggest that increased expression of MMR, aEJ, and BER pathway genes in *BRCA1*-and *BRCA2*-mutant breast tumors does not re-establish genomic stability. Instead, it correlates with higher FGA, implying that activation of these pathways represents adaptive dependencies that help maintain tumor viability by mitigating otherwise lethal DNA damage and allowing continued growth despite widespread genomic alterations. Moreover, the observed discrepancies between *BRCA1-* and *BRCA2*-mutant tumors indicate gene-specific utilization of shared repair pathways within distinct homologous recombination–deficient backgrounds. Together, these findings support a model in which MMR, aEJ, and BER pathway upregulation functions as a compensatory survival mechanism in BRCA-deficient tumors and represents an attractive class of therapeutic vulnerabilities.

### Breast tumors with *BRCA* deficiency is associated with adaptive engagement of replication stress tolerance networks

BRCA1 and BRCA2 maintain genome stability not only through their canonical HRR functions, but also by alleviating replication stress *via* multiple complementary mechanisms, including suppression of structural variation, protection of stalled replication forks from nuclease-mediated degradation, and restriction of ssDNA gap accumulation^60^. Despite experiencing substantial replication stress, established *BRCA*-mutant breast tumors often exhibit a striking ability to tolerate these conditions^61^. We hypothesized that *BRCA*-mutant tumors engage compensatory RST programs that preserve tumor fitness under persistent genomic stress. To test this, we systematically interrogated RST pathways and associated gene network dependencies in *BRCA*-mutant breast tumors.

Because *BRCA*-deficient cells fail to adequately restrain DNA replication under stress, replication fork reversal represents a key protective response that slows fork progression and limits fork collapse or degradation^62^. We assessed the expression of three established fork reversal factors HLTF, BLM, and FBH1^63–65^ and evaluated their relationship to FGA. In *BRCA1*-mutant tumors, high *HLTF* and *BLM* expression was associated with moderate-to-high FGA while tumors with low *HLTF* and *BLM* expression exhibited significantly lower FGA. *FBH1* showed a similar directional trend, although not statistically significance **(Fig. 2A)**. In *BRCA2-*mutant tumors, elevated *BLM* expression was likewise associated with increased FGA, supporting engagement of fork remodeling programs in the absence of BRCA2. However, neither *HLTF* nor *FBH1* expression showed a statistically significant relationship with FGA in this context **(Fig. S2A)**. Together, these results support a model in which *BRCA*-mutant breast tumors adaptively utilize RST mechanisms, with distinct helicase dependencies shaped by whether *BRCA1* or *BRCA2* is inactivated.

**Figure 2.**
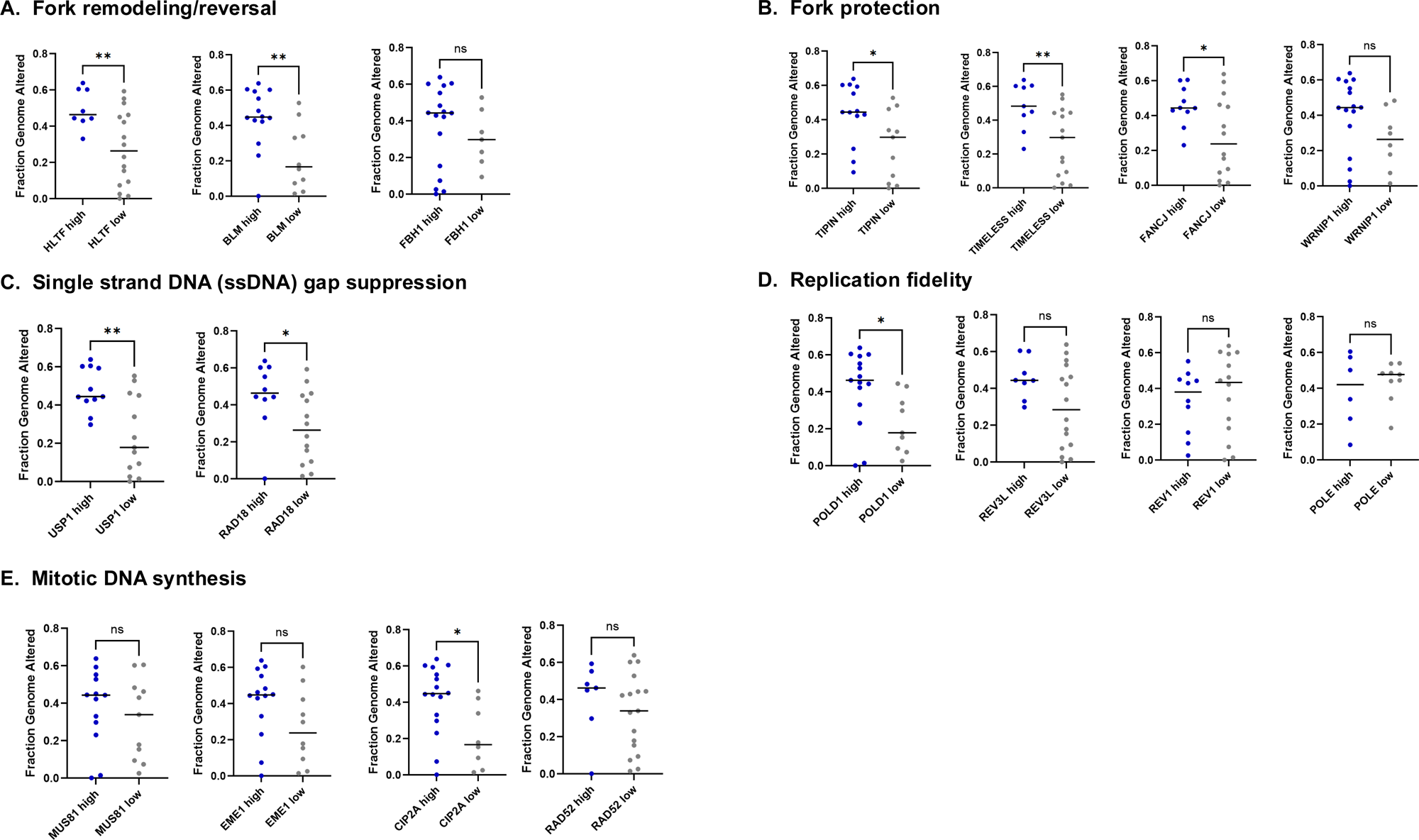
Replication stress tolerance pathways are selectively upregulated in *BRCA1* mutant breast tumors. FGA levels in TCGA *PanCancer Atlas* dataset (n = 24) for all genes shown except POLE (TCGA *Nature 2012*, n = 15) grouped into different stress tolerance mechanisms including **(A)** Fork remodeling/reversal, **(B)** Fork protection, **(C)** Single strand DNA (ssDNA) gap suppression**, (D)** Replication Fidelity or **(E)** Mitotic DNA synthesis. Low expression (heatmap mRNA expression z-score <0) and High expression (heatmap mRNA expression z-score >0). Asterisks denote significant p-value using two-tailed unpaired t-test with ***p<0.001, **p<0.01, *p<0.05; ns = non-significant.

To identify candidate pathways that may substitute for BRCA-dependent fork protection, we evaluated the expression of genes previously implicated in replication fork stabilization, focusing on *FANCD2, RAD51, TIPIN, TIMELESS, FANCJ,* and *WRNIP*. In *BRCA1*-mutant tumors, high expression of *FANCD2, RAD51, TIPIN, TIMELESS,* and *FANCJ* was associated with moderate-to-high FGA values **(Fig. 1B, 1C, and 2B)**. All these genes (except *FANCJ*) showed a similar directional relationship in *BRCA2*-mutant tumors; however, this association was only statistically significant for *TIPIN* **(Fig. S1B, S1C and Fig. S2B)**. These finding suggest that *BRCA*-mutant tumors partially compensate for defective fork protection through alternative fork-stabilizing programs, with distinctions emerging across *BRCA1*- and *BRCA2*-mutant contexts.

Cells can engage an alternative RST pathway mediated by PRIMPOL which promotes replication fork restart by repriming DNA synthesis downstream of lesions; however, this mechanism leaves behind ssDNA gaps, representing a distinct and potentially deleterious form of replication-associated DNA damage^66–68^. USP1 limits the accumulation of these ssDNA gaps by regulating post-replication repair, including de-ubiquitination of key substrates such as PCNA^69,70^. RAD18 can similarly limit gap-associated damage by promoting PCNA polyubiquitination to activate template switching (an error-free gap repair pathway) and by facilitating RAD51 loading at post-replicative ssDNA gaps^71,72^. We investigated the correlation between *USP1* expression and FGA in *BRCA1-* and *BRCA2-*mutant cancers. In *BRCA1-*mutant tumors, high *USP1* and *RAD18* expression significantly correlated with moderate-to-high FGA values **(Fig. 2C)**, suggesting that elevated activity of these pathways may enable *BRCA1*-mutant tumors to tolerate heightened replication stress and chromosomal instability. *BRCA2*-mutant tumors showed a similar directional trend, albeit not significant, potentially reflecting *BRCA* genotype–specific biology or limited statistical power in this cohort **(Fig. S2C)**

POLD1 (DNA polymerase δ1) is a key replicative polymerase with both polymerase activity and 3′–5′ exonuclease proofreading function, particularly essential for DNA replication on the lagging strand^73,74^.POLD1 expression has been associated with aggressive disease and advanced stage in different cancer types ^75,76^. We stratified *BRCA1*- and *BRCA2*-mutant tumors based on *POLD1* expression, and found that high *POLD1* expression was associated with moderate-to-high FGA values in both *BRCA1-* and *BRCA2-*mutant cohorts **(Fig. 2D and Fig. S2D)**. This suggests that POLD1 stabilizes replication forks and supports continued proliferation despite genomic instability in BRCA-deficient tumors, thereby reflecting an adaptive response of breast cancer cells to genomic stress.

Translesion synthesis (TLS) polymerases facilitate continued DNA replication across damaged templates, thereby preventing sustained replication-fork stalling. However, this tolerance mechanism typically compromises replication fidelity because TLS polymerases lack the robust proofreading capacity of canonical replicative polymerases^77,78^. We interrogated TLS-associated gene networks with a focus on REV3L, REV1, and POLE. REV1 primarily functions as a scaffold that coordinates recruitment of TLS polymerases to sites of damage, whereas REV3L encodes the catalytic subunit of DNA polymerase ζ (Pol ζ), the principal extender polymerase that promotes lesion bypass. POLE encodes the catalytic subunit of DNA polymerase ε (Pol ε), the major leading-strand replicative polymerase that also contributes to genome maintenance and DNA damage responses^79,80^. We investigated FGA levels in *BRCA1*-mutant tumors stratified by *REV3L, REV1,* and *POLE* expression. High *REV3L* expression was associated with a modest increase in FGA, consistent with a trend toward greater genomic instability in tumors with elevated Pol ζ expression, although this difference did not reach statistical significance **(Fig. 2D and S2D)**. In contrast, variation in *REV1* or *POLE* expression showed no clear relationship with FGA **(Fig. 2D and S2D)**, indicating that TLS components do not contribute uniformly to genome-wide instability. These findings suggest that increased reliance on specific TLS factors, particularly REV3L, may confer a limited survival advantage in *BRCA1/2*-mutant breast tumors.

Because BRCA-deficient cells cannot perform HR, they depend on alternative mitotic DNA repair pathways to survive replication stress^81,82^. We therefore examined alternative mitotic repair programs in *BRCA1/2*-mutant breast tumors, focusing on MUS81, EME1, CIP2A, and RAD52. MUS81–EME1 is a structure-specific endonuclease that resolves abnormal DNA intermediates, including Holliday junction–like structures and stalled or under-replicated forks, a function especially important in mitosis for accurate chromosome segregation^83,84^. CIP2A serves as a scaffold that recruits the SLX4–MUS81–XPF nuclease complex during mitosis to process persistent replication-associated DNA structures^85^. RAD52 supports this pathway by promoting MUS81 recruitment and enabling the break-induced replication (BIR)-like component of mitotic DNA synthesis (MiDAS)^86,87^. In the investigated TCGA *PanCancer Atlas* study, we found that higher expression of these genes showed a general trend toward increased FGA in *BRCA1-* and *BRCA2-*mutant tumors, consistent with a modest association between elevated MiDAS-related activity and genomic instability; however, statistical significance was observed only for *CIP2A* in *BRCA1-*mutant tumors and for EME1 in *BRCA2-mutant* tumors **(Fig. 2E and Fig. S2E)**. In the *Nature 2012* cohort, *BRCA2-*mutant tumors with high *RAD52* expression exhibited significantly higher FGA **(Fig. S2E)**. There is therefore a selective upregulation of specific MiDAS components which may modestly reduce replication-associated damage, although the overall effect appears limited. To determine whether prolonged FANCM depletion promotes progressive genome instability in BRCA1-deficient breast cancer cells, we depleted FANCM in cancer cell line and found that the prolong loss of FACNM induces newly acquired tandem duplications, mainly class 1 **(Fig. S2F)**. These analyses indicate that *BRCA1/2*-mutant breast tumors engage multiple adaptive RST pathways that vary by *BRCA* genotype.

### Tumor grade-dependent enrichment of DNA repair and replication stress tolerance programs

After identifying potential compensatory DDR and RST pathways wherein high FGA correlated with an upregulation in expression of their associated genes, we asked whether these dependencies are preferentially enriched in specific tumor grades. Tumor grade was evaluated relative to primary tumor stage (T1–T4), where higher T stages correspond to increased tumor size and more extensive local tissue invasion^88^.

Among breast tumors with high HRR genes *DNA2* and *RAD51C* expression, T2 and T4 tumors exhibited significantly greater FGA than T1 tumors. A similar pattern was observed in tumors with elevated *EXO1, RAD51,* and *BARD1* expression, but the increase in FGA was statistically significant in T2 compared with T1 tumors **(Fig. 3A).** For MMR, tumors with increased *MSH3* expression showed significantly elevated FGA in T2, T3 and T4 relative to T1 tumors. Tumors with elevated *MSH2* and *MSH6* expression also demonstrated a trend toward higher FGA with increasing grade; however, statistical significance was observed only for T2 versus T1 in the *MSH2*-high subgroup **(Fig. 3B).** For FA pathway, T2 and T4 tumors with elevated *FANCM* expression displayed significantly higher genome instability than T1 tumors. Likewise, among tumors with increased *FANCE* and *FANCD2* expression, FGA was significantly increased in T2 tumors compared with T1 **(Fig. 3C)**. In contrast, there was no significant difference in FGA across all stages for tumors with high *FANCA, B* or *C* expression. Focusing on BER-associated genes, tumors with high *APE2* expression were associated with significantly elevated genomic instability in T2 tumors; this grade-associated pattern was not observed for tumors with high *FEN1* or *UNG2* expression **(Fig. 3D).** Finally, in the alternative end-joining (aEJ) pathway, tumors with high *XRCC1* expression exhibited significantly increased FGA in T2 compared with T1, while there was no significant difference observed in POLQ-high tumors **(Fig. 3E)**

**Figure 3.**
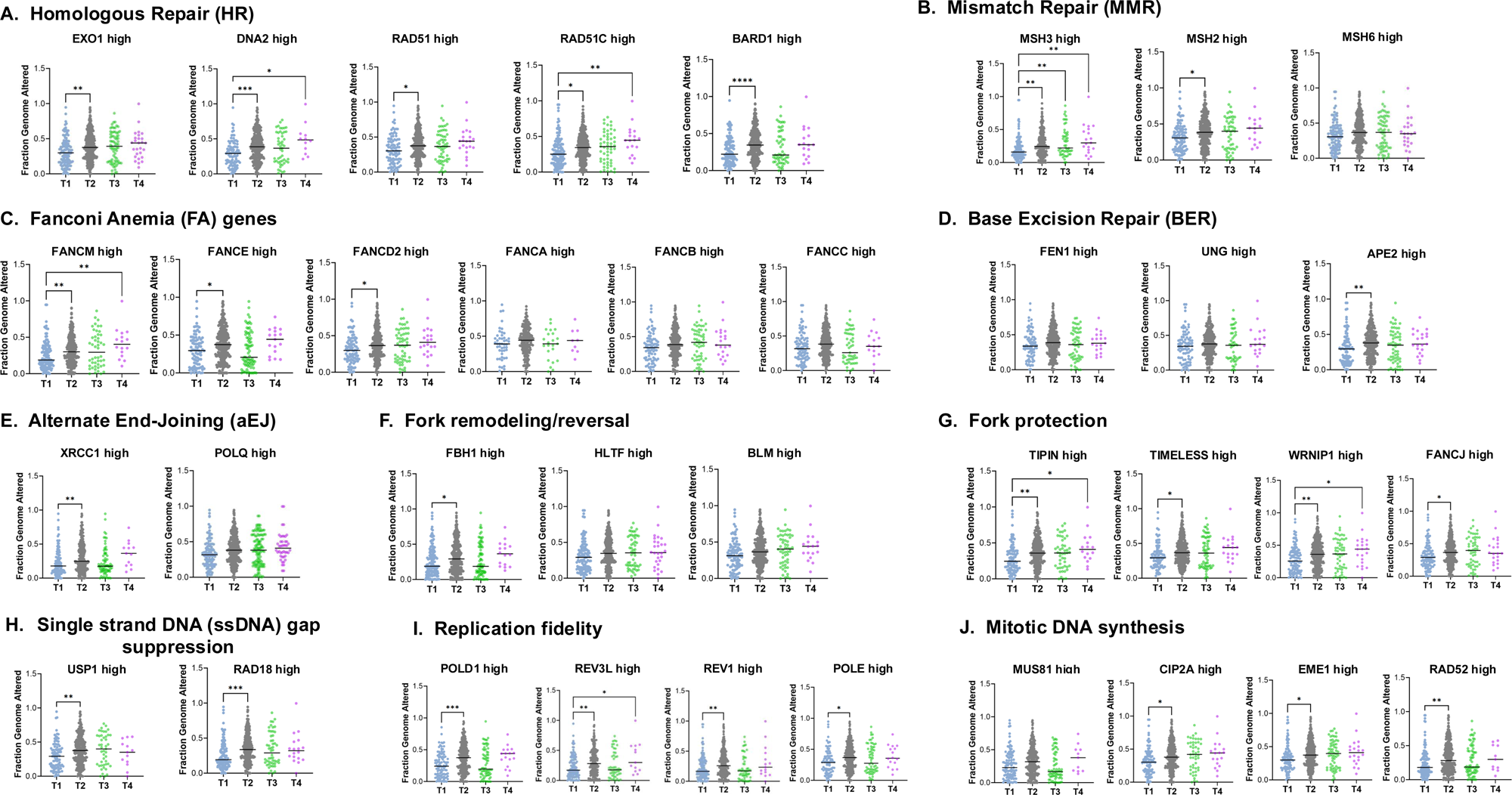
Progressive tumor grades are associated with increased reliance on DNA repair and RST pathways. FGA levels across 4 tumor stage codes developed by the American Joint Committee on Cancer (AJCC)—T1, T2, T3, and T4, where T refers to tumor size as classified by the TNM staging system (Tumor, Node, Metastasis), using TCGA *PanCancer Atlas* dataset for breast invasive carcinoma, irrespective of *BRCA1/2* mutation status. For each graph, only patients having high expression of indicated genes (heatmap mRNA expression z-score > 0) were selected and genes were grouped into the different DNA-repair pathways including (A) Homologous repair, (B) Mismatch repair, (C) Fanconi Anemia (FA), (D) Base excision repair (BER), (E) Alternative end-joining; or stress tolerance mechanisms including (F) Fork remodeling, (G) Fork protection, (H) single strand DNA gap suppression, (I) replication fidelity or (J) mitotic DNA synthesis mechanisms. Asterisks denote significant p-value using Tukey’s multiple comparisons test with ****p<0.0001, ***p<0.001, **p<0.01, *p<0.05.

We then asked whether replication stress response and tolerance programs exhibit similar grade-dependent enrichment. Among fork remodeling genes, high FBH1 expression was associated with significantly increased genomic instability in T2 relative to T1 tumors, whereas no significant grade-associated differences were detected for HLTF or BLM **(Fig. 3F)**. In contrast, high FGA showed a consistent and significant association with T2 tumors having high expression of fork protection genes *TIPIN, TIMELESS, WRNIP*, or *FANCJ* relative to T1 tumors, and *TIPIN-* and *WRNIP*-high tumors also showed significantly higher FGA in T4 *versus* T1 disease **(Fig. 3G)**. For ssDNA gap suppression, tumors with elevated *USP1* or *RAD18* expression displayed a very significant increase in FGA in T2 compared with T1 **(Fig. 3H)**. Similar significant associations were observed for replication fidelity genes *POLD1, REV3L, REV1,* and *POLE* **(Fig. 3I)**. Finally, within the mitotic DNA synthesis pathway, high *CIP2A, EME1*, and *RAD5*2 expression was associated with significantly increased FGA in T2 tumors compared to T1, whereas *MUS81*-high tumors did not show a statistically significant grade-dependent effect **(Fig. 3J)**

Taken together, these results indicate that as breast tumors advance to more locally aggressive stages, they increasingly depend on the upregulation of different DNA repair and RST pathway as an adaptive strategy.

### Adaptive genome-maintenance programs are preferentially associated with Basal-like and Luminal-B subtypes

We asked whether compensatory DNA repair and RST pathways–associated genes are preferentially enriched within specific molecular subtypes. To address this, patients with high expression of adaptive-dependency–associated genes were stratified by subtype, and genomic instability was quantified using FGA across four clinically relevant subtypes: Luminal A, Luminal B, HER2-enriched, and basal-like (TNBC).

Across pathways, elevated expression of repair and RST genes consistently delineated basal-like tumors as the most genomically unstable subtype, with Luminal B typically exhibiting intermediate instability relative to Luminal A. Within HRR, tumors with high *EXO1, DNA2, RAD51, RAD51C*, or *BARD1* expression displayed significantly higher FGA in basal-like than in Luminal A and HER2-enriched tumors; in these same high-expression strata, Luminal B tumors also showed significantly greater genomic instability than Luminal A **(Fig. 4A)**. MMR analyses further reinforced this pattern: in tumors with elevated *MSH2* or *MSH6* expression, basal-like tumors exhibited significantly greater genomic instability than Luminal A and HER2-enriched tumors **(Fig. 4B)**. Elevated *MSH3* expression similarly stratified basal-like tumors with higher FGA relative to Luminal A, although the comparison with HER2-enriched tumors did not reach statistical significance **(Fig. 4B)**. Across all MMR-high groups, Luminal B tumors also demonstrated significantly higher FGA than Luminal A. A similar subtype hierarchy was observed for the FA pathway: among tumors with elevated *FANCD2, FANCB, FANCC, FANCE, FANCA*, or *FANCM* expression, basal-like tumors exhibited significantly higher FGA than Luminal A, and—specifically for *FANCD2, FANCB, FANCC*, and *FANCE*—also higher FGA than HER2-enriched tumors **(Fig. 4C)**. Moreover, in most FA-high subgroups (all except *FANCA*), Luminal B tumors again showed significantly increased FGA compared with Luminal A **(Fig. 4C)**. We next evaluated additional repair pathways: in BER, tumors with elevated *FEN1, UNG,* or *APE2* expression displayed significantly higher FGA in basal-like tumors compared with Luminal A, and Luminal B tumors again exhibited significantly greater instability than Luminal A **(Fig. 4D)**. Among aEJ-associated *POLQ*-high and *XRCC1*-high tumors, basal-like disease exhibited significantly higher FGA than Luminal A and HER2-enriched tumors, which was also significantly higher compared to Luminal B tumors with high *XRCC1* expression **(Fig. 4E)**. Additionally, in *POLQ*-high tumors Luminal B showed higher instability than Luminal A; in *XRCC1*-high tumors, both Luminal B and HER2 tumors exhibited significantly higher genome instability compared to Luminal A tumors **(Fig. 4E)**.

**Figure 4.**
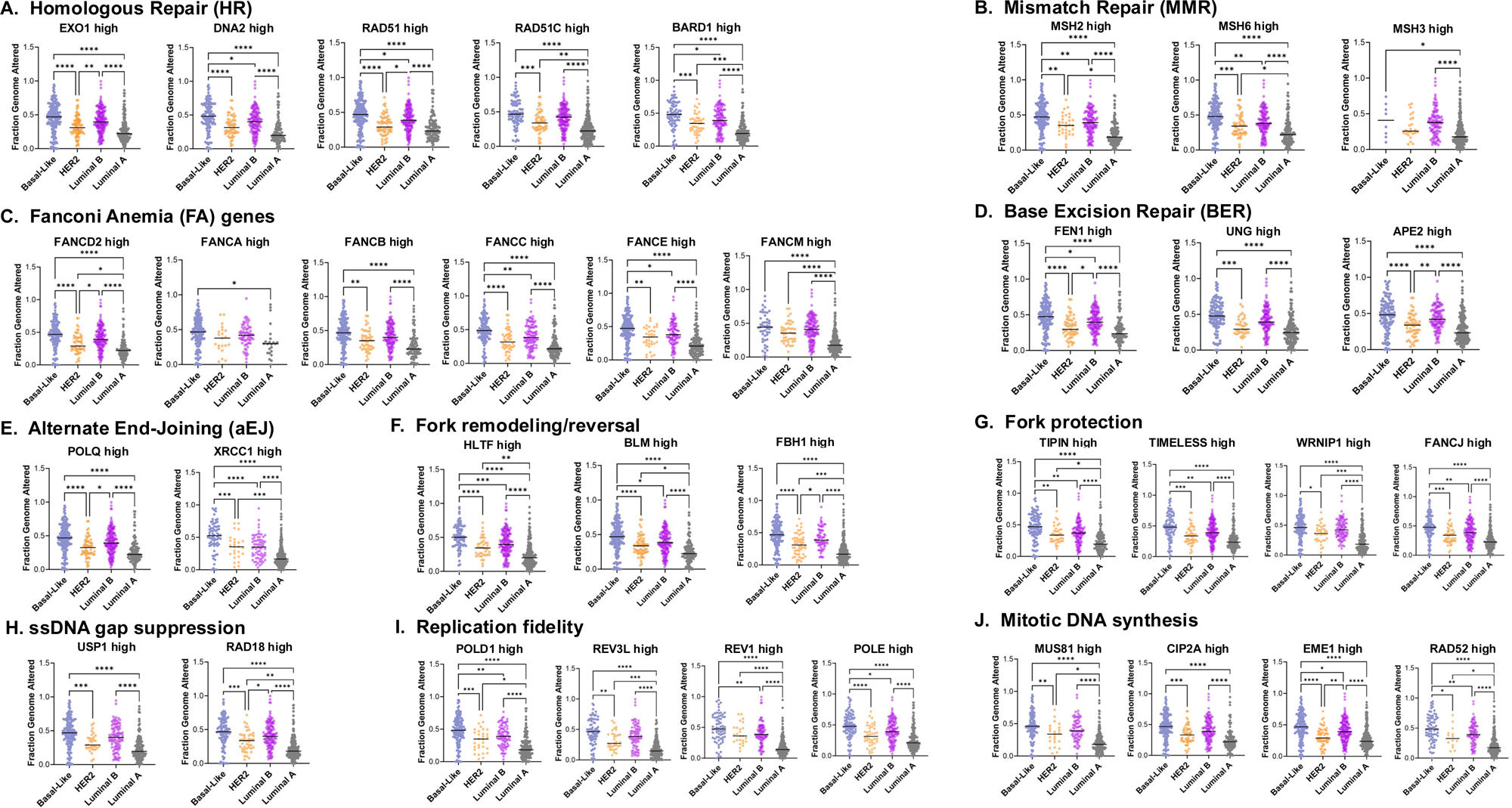
Basal-like and luminal B tumors show enhanced reliance on adaptive genome maintenance programs. FGA levels across 4 different breast cancer subtypes organized from most to least aggressive: Basal-Like, HER2, Luminal B and Luminal A, using the TCGA *PanCancer Atlas* dataset for breast invasive carcinoma irrespective of *BRCA1/2* mutation status. For each graph, only patients having high expression of indicated genes (heatmap mRNA expression z-score >0) were selected and genes were grouped into the different DNA-repair including (A) Homologous repair, (B) Mismatch repair, (C) Fanconi Anemia (FA), (D) Base excision repair (BER), (E) Alternative end-joining; or stress tolerance mechanisms including (F) Fork remodeling, (G) Fork protection, (H) single strand DNA gap suppression, (I) replication fidelity or (J) mitotic DNA synthesis mechanisms. Asterisks denote significant p-value using Tukey’s multiple comparisons test with ****p<0.0001, ***p<0.001, **p<0.01, *p<0.05.

We then asked whether RST programs show a comparable subtype enrichment. In tumors with elevated expression of fork remodeling genes (*HLTF, BLM,* and *FBH1*), the basal-like subtype exhibited significantly higher FGA than Luminal A, Luminal B (except for *FBH1*), and HER2-enriched tumors **(Fig. 4F)**. Further, Luminal B tumors showed significantly higher FGA than Luminal A tumors **(Fig. 4F)**. Fork protection programs followed a similar pattern: tumors with high *TIPIN, TIMELESS, WRNIP1*, or *FANCJ* expression showed significantly higher FGA in basal-like tumors than in Luminal A and Luminal B tumors were significantly more unstable than Luminal A **(Fig. 4G)**. Comparable subtype stratification was observed for ssDNA gap suppression (*USP1, RAD18*) and replication fidelity (*POLD1, REV3L, REV1, POLE*), where basal-like tumors exhibited significantly higher FGA than Luminal A (and, in some comparisons, Luminal B), and Luminal B remained significantly more unstable than Luminal A **(Fig. 4H and 4I)**. Finally, within the mitotic DNA synthesis pathway, tumors with elevated *MUS81, CIP2A, EME1*, or *RAD52* expression showed significantly increased FGA in basal-like tumors compared with Luminal A, with Luminal B again exhibiting significantly higher FGA than Luminal A **(Fig. 4J)**.

Collectively, these results demonstrate that elevated expression of multiple DNA repair and RST programs preferentially marks basal-like—and, to a lesser extent, Luminal B—breast tumors with the highest genomic instability, consistent with subtype-specific adaptive reliance on genome-maintenance pathways to tolerate an increased genome alteration burden.

### Breast tumor mutational burden (TMB) is a pathway specific consequence of disrupted DNA repair and replication stress tolerance

Tumor mutational burden (TMB) is a clinically relevant genomic biomarker associated with responsiveness to immune checkpoint inhibitors (ICIs) across multiple cancer types^36^. To test whether elevated TMB is a generalized consequence of genomic instability or selective outcome of specific DDR and RST defects, we analyzed the TCGA *PanCancer Atlas* study, with samples stratified by pathway and gene-level status across canonical DNA repair, fork protection and replication fidelity programs, and quantified mutation burden. We used mutation count as a metric, defined here as the total number of somatic point mutations including single-nucleotide variants and small insertions/deletions (indels).

Within HR, increases in mutation burden were comparatively selective. Mutation count in patients harboring *BRCA2* mutation was significantly relative to total mutation count, whereas other HR factors showed smaller or non-significant shifts **(Fig. 5A)**. This pattern suggests that only a subset of HR-associated dependencies track tightly with global mutational load. The most pronounced mutation count was observed for MMR genes. Tumors with mutations in *MSH2, MSH3*, or *MSH6* showed robust, highly significant increases in mutation counts relative to “All” **(Fig. 5A)**. Importantly, the consistency across multiple MMR components suggests that this association reflects pathway-level biology rather than idiosyncratic single-gene effects. Elevated mutation burden was also observed with alternative end joining (aEJ), with patients harboring *POLQ* or *XRCC1* mutations showing significantly higher mutation counts. This is consistent with aEJ activity being coupled to error-prone repair and mutation accumulation under conditions of elevated DNA damage or replication stress **(Fig. 5A)**.

**Figure 5.**
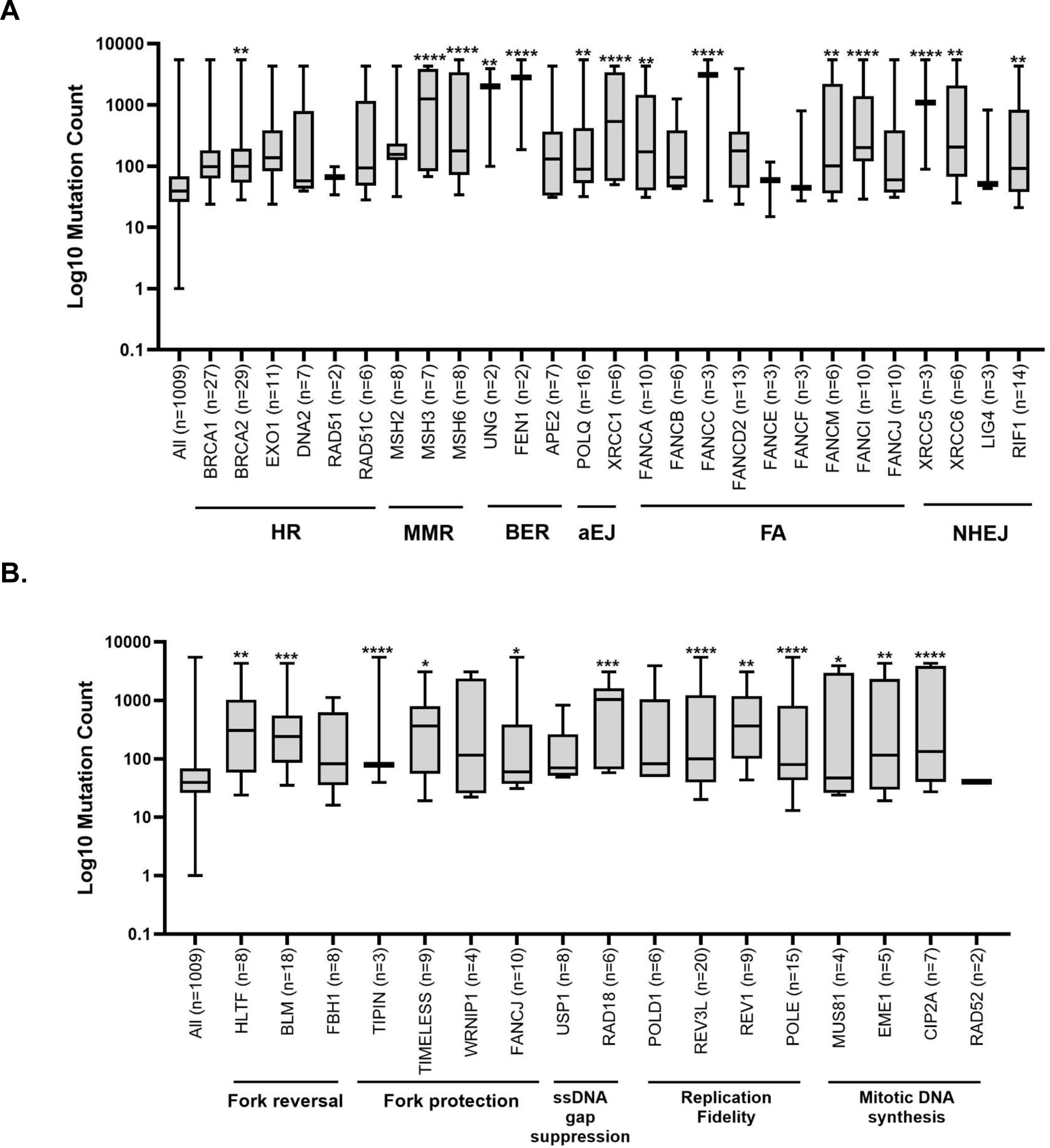
Tumor mutation burden is selectively enriched in discrete DNA repair and replication stress pathway defects. Box-and-whisker plots showing median, interquartile range, and minimum–maximum values of Log10 mutation count in samples containing mutations within specific genes involved in different **(A)** DNA repair pathways and **(B)** stress tolerance mechanisms analyzed in TCGA *PanCancer Atlas* excluding those with mutation count = 0 (n= 1009). The number of samples with mutations in each particular gene is indicated between parenthesis. Asterisks denote significant p-value using Tukey’s multiple comparisons test with ****p<0.0001, ***p<0.001, **p<0.01, *p<0.05.

FA pathway-associated gene *FANCA* and *FANCC* demonstrated a clearer association with increased mutation burden showing significant increases in mutation burden relative to “All”, with *FANCC* exhibiting one of the strongest effects among FA components **(Fig. 5A)**. Other FA genes displayed broader distributions without consistent significance, indicating heterogeneity in the extent to which FA-associated gene strata capture hypermutated tumors. BER genes, namely *UNG* and *FEN1*, showed significant enrichment for increased mutation burden. *XRCC5* and *XRCC6* also showed significant increases **(Fig. 5A)**. By comparison, canonical NHEJ factors showed weaker or absent associations with mutation burden. For example, *LIG4* did not exhibit a significant increase relative to “All”, whereas *RIF1* showed a more modest but significant elevation **(Fig. 5A)**. Together, these results underscore pathway specificity in the relationship between DDR gene and hypermutation.

We next evaluated whether disruption of RST and replication fidelity programs similarly partitions tumors by mutation burden. Within fork reversal, mutation count was significantly elevated in *HLTF*- and *BLM*-stratified tumors, whereas *FBH1* did not reach significance **(Fig. 5B)**. Fork protection defects also associated with increased mutation burden, with significant elevations in *TIPIN*-, *TIMELESS*-, and *FANCJ*-stratified subsets, while *WRNIP1* did not reach significance. Defects in ssDNA gap suppression showed a strong association with elevated mutation burden in *RAD18*-stratified tumors, whereas *USP1* did not reach significance. Replication fidelity defects were likewise linked to increased mutation burden, with *POLD1-, POLE-, REV3L-,* and *REV1*-stratified subsets all showing highly significant elevations relative to “All”. Finally, within mitotic DNA synthesis, mutation burden was significantly increased in *MUS81-, EME1*-, and *CIP2A*-stratified tumors, whereas *RAD52* did not reach significance **(Fig. 5B)**. Collectively, these analyses reveal that TMB is not uniformly distributed across disrupted genome maintenance processes; instead, it concentrates within discrete DDR/RST states.

### Co-occurrence and mutual exclusivity map heterogeneous repair pathway engagement

Cooperative genetic interactions, reflected by either patterns of co-occurrence or mutually exclusive relationships, shape genomic instability, therapeutic responsiveness, and immune engagement, thereby informing the rational design of combination therapies and precision medicine^89–93^. We focused on these two classes of genetic interactions between the six most frequently mutated driver genes (*TP53, BRCA1, BRCA2, PIK3CA, FANCD2 and BAP1*) in breast cancer and genes involved in DNA repair (HR, MMR or BER) ^94,95^. As expected, focusing on basal-like breast cancer samples (n= 171) in the TCGA *PanCancer Atlas* study, *TP53* mutations dominated the cohort, occurring in almost 90% of samples. Other recurrent driver alterations were observed at substantially lower frequencies (<10%), including *BRCA1* (7.6%), *PIK3CA* (7.0%), *BRCA2* (3.5%), *FANCD2* (3.5%), and *BAP1* (2.3%) **(Fig. 6A)**. This distribution underscores the central role of *TP53* disruption in basal-like disease, with additional drivers contributing to smaller subsets of tumors.

**Figure 6.**
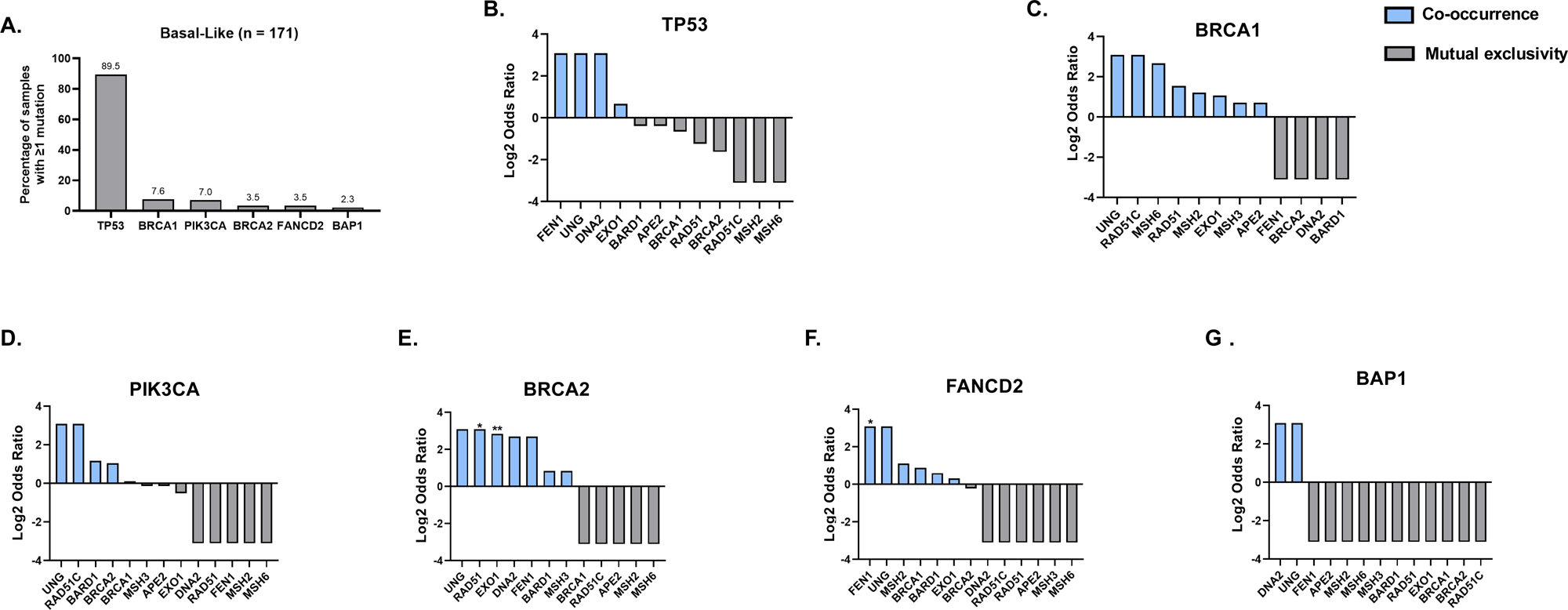
Co-occurrence and mutual exclusivity reveal heterogeneous DNA repair dependencies in basal-like breast cancer. **(A)** Percentage of patients with mutation in the top mutated genes in breast cancer focusing on Basal-Like samples (n = 171) from the TCGA *PanCancer Atlas* dataset. Tendency (Log2 Odds Ratio) toward co-occurrence (positive values; blue bars) or mutual exclusivity (negative values; grey bars) for **(B)** *TP53*, **(C)** *BRCA1*, **(D)** *PIK3CA*, **(E)** *BRCA2*, **(F)** *FANCD2*, and **(G)** *BAP1* with genes involved in HR, MMR or BER (as described in previous figures). Asterisks denote significant p-value **p<0.01, *p<0.05 using two-sided Fisher’s Exact Test.

Pairwise interaction analyses revealed distinct and gene-specific patterns of co-occurrence and mutual exclusivity across DNA damage response pathways. *TP53* alterations preferentially co-occurred with BER factors *FEN1* and *UNG* and HR factors *DNA2* and *EXO1*, while exhibiting mutual exclusivity with MMR genes *MSH2* and *MSH6*, as well as HR genes *BRCA1*, *BRCA2*, and *RAD51C* **(Fig. 6B)**. These patterns suggest complementary roles between *TP53* loss and BER activity, contrasted with functional redundancy or incompatibility with MMR and HR defects. Tumors harboring *BRCA1* mutations display co-occurrence with *RAD51C* and MMR genes, including *EXO1* and *MSH6*, but were mutually exclusive with long-patch BER components such as *FEN1* and *DNA2* **(Fig. 6C)**. In contrast, *PIK3CA-*mutant tumors showed co-occurrence with *UNG* and *RAD51C*, alongside a tendency toward mutual exclusivity with MMR genes *MSH2* and *MSH6*, and BER factors *FEN1* and *DNA2*, indicating a distinct pattern of pathway engagement **(Fig. 6D)**.

*BRCA2*-mutant tumors demonstrated significant co-occurrence with core HR components, including *RAD51* and *EXO1*.They also showed a tendency of co-occurrence with BER factors *FEN1* and *UNG*, while remaining mutually exclusive with MMR genes *MSH2* and *MSH6* **(Fig. 6E)**. *FANCD2* alterations similarly preferentially co-occurred with BER factors *FEN1*, which was significant, and *UNG*, but were largely mutually exclusive with HR genes *RAD51* and *RAD51C* and with MMR genes *MSH3* and *MSH6* **(Fig. 6F)**. In contrast to these more selective interaction patterns, *BAP1* mutations exhibited broad mutual exclusivity with the selected DNA repair genes of interest across the DNA damage response genes examined, with the exception of *DNA2* and *UNG*, suggesting a distinct and potentially independent role for *BAP1* loss in basal-like breast cancer **(Fig. 6G)**. Together, these results demonstrate that basal-like breast cancers are organized into distinct genetic interaction states defined by non-random patterns of co-occurrence and mutual exclusivity among DNA damage response pathways, revealing heterogeneous repair dependencies that may be exploited for precision therapeutic stratification.

## DISCUSSION

This study identifies an integrated model in which genomic instability in BC is not merely tolerated but actively managed through adaptive rewiring of DDR and RST networks. This concept aligns with prior work describing “non-oncogene addiction” to stress response pathways, where tumor cells rely on DDR signaling for survival under intrinsic stress^96,97^. Our findings extend these models by resolving this adaptation at the pathway level, demonstrating that genomic instability is coupled to selective, rather than global, activation of DDR and RST programs. This refines earlier models of HRD, which often treat genome instability as a binary state^14,98^, by showing that tumors actively reconfigure multiple repair and tolerance pathways in a context-dependent manner.

By integrating genomic instability metrics, gene expression dependencies, tumor grade, molecular subtype, mutational burden, and genetic interaction mapping, our findings converge on a central concept: breast tumors persist not by restoring genome integrity, but by engaging compensatory, often error-prone, genome-maintenance programs that permit survival under chronic genotoxic stress. We distinguish between repair restoration and damage tolerance is consistent with prior observations in *BRCA*-mutant systems, where partial HR activity or reversion does not fully normalize genome stability^99,100^. However, our data suggest a broader and more systematic phenomenon, in which multiple pathways—not just HR—contribute to buffering replication-associated damage. If compensatory pathway activation re-established canonical repair, one would predict reduced genome alteration in high-expression groups. Instead, we observed the opposite pattern across many gene strata in *BRCA*-mutant tumors, consistent with adaptive buffering that preserves viability under chronic genotoxic stress while permitting continued genomic evolution. This interpretation provides a mechanistic framework for the clinical paradox in *BRCA*-mutant BC: profound genomic instability coexisting with sustained tumor fitness and progression^101,102^.

Across independent cohorts, *BRCA1*-mutant tumors showed consistently higher FGA, and *BRCA2-*mutant tumors showed a similar directional trend, supporting the central premise—well established in both experimental models and clinical sequencing studies—that BRCA loss drives chromosomal instability^103,104^. Within this context, increased expression of specific homologous recombination, Fanconi anemia, mismatch repair, alternative end joining, and base excision repair factors associated with higher genome alteration. This contrasts with classical models in which upregulation of repair genes is interpreted as restoration of genomic integrity, and instead supports emerging literature suggesting that overexpression of repair components can promote aberrant or alternative repair outcomes as a protective survival mechanism^105,106^.

Notably, HR-associated components such as EXO1, DNA2, RAD51, and RAD51C remain transcriptionally active in *BRCA1/2*-mutant tumors and associate with higher FGA. Prior studies have shown RAD51 loading and resection activity can persist in *BRCA*-deficient contexts, but often lead to incomplete or error-prone repair^107,108^; our findings extend this by linking such activity directly to increased chromosomal alteration in clinical tumors. Similarly, FA pathway activation, traditionally linked to ICL repair, tracks with increased genomic instability, consistent with reports that FA components promote replication fork stability and restart rather than prevent damage accumulation^18,109^. Importantly, the lack of significant association between NHEJ components and FGA contrasts with some earlier assumptions that NHEJ dominates repair in HR-deficient tumors^110^, and instead highlights pathway specificity in compensatory rewiring.

Beyond DNA repair, our results identify RST pathways as a critical layer of adaptation in *BRCA*-mutant breast tumors. This is consistent with a growing body of literature positioning replication stress as a central driver of tumor evolution, particularly in oncogene-activated and HR-deficient cancers^111,112^. Elevated expression of fork remodeling (HLTF, BLM), fork protection (FANCD2, TIPIN, TIMELESS), post-replicative gap suppression (USP1, RAD18), and replication fidelity (POLD1) factors consistently associates with higher genomic instability. Prior mechanistic studies have demonstrated that fork protection and remodeling factors prevent catastrophic fork collapse, but can also enable survival with under-replicated or misrepaired DNA^19,113^. Our findings provide clinical-scale evidence that these processes are not merely protective but are actively selected in highly unstable tumors. The relatively modest contribution of TLS factors, with only REV3L showing a weak association with FGA, is notable given prior emphasis on translesion synthesis in damage tolerance^114^, and suggests that fork-centric mechanisms may play a more dominant role than previously appreciated in BC.

Our data further indicate that compensatory dependencies are non-uniform and genotype-specific, even within the same pathway. This observation is consistent with recent work demonstrating that BRCA1 and BRCA2 deficiencies generate distinct replication-associated lesions and repair liabilities^20,115^. For example, *BRCA1-* and *BRCA2-*mutant tumors shared some directional patterns but diverged in statistical enrichment for multiple RST nodes, including fork remodeling, fork protection, gap suppression, and mitotic DNA synthesis components. This heterogeneity reinforces emerging critiques of the “BRCAness” concept as overly reductive^14,17^, and supports a model in which HRD represents a spectrum of mechanistically distinct states.

Our analyses across tumor grades and molecular subtypes reveal that adaptive genome-maintenance programs are not static but evolve with disease progression, aligning with longitudinal and evolutionary studies showing that genomic instability and replication stress increase with tumor progression and therapeutic pressure^116,117^. As tumors progressed from T1 to higher local stages, expression-defined DDR/RST dependencies increasingly associate with elevated FGA, indicating that adaptation intensifies during progression. This dependency is especially pronounced in basal-like and Luminal B subtypes. Basal-like tumors, which overlap substantially with triple-negative breast cancer, have long been characterized by high genomic instability and replication stress signatures^47,118^; our findings provide mechanistic resolution by identifying the specific pathways supporting this state. These patterns also suggest that adaptive genome-maintenance programs are not merely passive markers of instability; rather, they likely represent active states selected during tumor evolution in aggressive clinical contexts.

Our mutational burden analyses extend this framework to tumor immunogenic potential. Consistent with extensive prior literature, mismatch repair deficiency shows the strongest association with elevated mutation burden^119,120^. However, our data refine this paradigm by demonstrating that not all forms of genomic instability contribute equally to TMB. While *BRCA2* mutations show only modest effects on mutation counts, alterations in aEJ, BER, and replication fidelity genes (e.g., POLQ, XRCC1, POLD1, POLE) show strong associations with increased TMB, consistent with known mutator phenotypes linked to polymerase proofreading defects^121,122^. In contrast, many HR and FA components do not show consistent associations with TMB, reinforcing the idea that chromosomal instability (captured by FGA) and point mutation burden (TMB) arise from distinct but overlapping biological processes^101^.

The co-occurrence and mutual exclusivity analyses further support the existence of structured, non-random repair pathway engagement. Patterns such as TP53 co-occurrence with BER factors and mutual exclusivity with MMR and HR genes are consistent with prior pan-cancer analyses showing that tumors avoid combinations of defects that would be synthetically lethal^123,124^. Mutual exclusivity between *BRCA1/2* and MMR components is particularly notable and aligns with experimental evidence that combined loss of these pathways is poorly tolerated. Conversely, co-occurring alterations may reflect cooperative mechanisms that enhance tumor fitness and contribute to resistance.

This study is based on integrative analysis of large-scale genomic/transcriptomic datasets, and therefore has inherent limitations. Gene expression does not always directly reflect pathway activity, and functional validation will be required to confirm the causal role of identified dependencies. Sample size constraints for some mutation-defined strata, especially *BRCA2*-mutant subsets, likely reduced statistical power for selected comparisons. In addition, mutation count was used as the TMB metric in this study; while informative for relative stratification, harmonization with standardized nonsynonymous-megabase TMB definitions and external immunotherapy-treated cohorts will be important for clinical translation. Despite these constraints, the consistency of pathway-level trends across independent cohorts and across complementary analyses (FGA, mutation burden, grade/subtype enrichment, and genetic interaction structure) supports a robust biological signal. Future work should focus on (i) Functional interrogation of candidate compensatory pathways in model systems; (ii) Longitudinal studies to track evolution of repair dependencies during treatment and (iii) Clinical validation of combination strategies targeting DDR/RST networks.

In summary, this study demonstrates that breast tumors, particularly those with BRCA deficiency, survive pervasive genomic instability through coordinated and heterogeneous engagement of compensatory DDR and RST pathways. These adaptive programs do not restore genome integrity but instead enable tolerance of ongoing damage, thereby sustaining tumor growth. By mapping these dependencies across tumor contexts, our work provides a framework for identifying actionable vulnerabilities and designing rational, combination-based therapeutic strategies to overcome resistance and improve patient outcomes.

## METHODS

### Fraction Genome Altered Relative to *BRCA* Gene Status

Publicly available datasets from three independent breast cancer cohorts were obtained via the cBioPortal platform (https://www.cbioportal.org): *TCGA PanCancer Atlas*, *MSK 2025*, and *TCGA Nature 2012*. All datasets were accessed under the “Breast” category, specifically within “*Invasive Breast Carcinoma”* studies. For each dataset, somatic mutations and putative copy-number alterations from GISTIC were selected for all subsequent analysis were selected for analysis. Queries were performed using either *BRCA1* or *BRCA2* as the gene of interest, and all available samples were included. Samples were stratified based on gene status into *mutant* and *wild-type* groups. Cases lacking mutation profiling data for *BRCA1/2* were filtered out. Fraction of Genome Altered (FGA) was extracted as a clinical attribute for each sample. Corresponding patient-level data, including patient identifiers, *BRCA1/2* mutation status, and FGA values, were downloaded for downstream analysis. All statistical analyses and data visualization were performed using GraphPad Prism (version 10.6.1). Differences in FGA between *BRCA*-mutant and wild-type groups were assessed for statistical significance using appropriate tests as indicated in the figure legends. Sample sizes are reported in the respective figure legends.

### Gene Expression Patterns Relative to FGA in BRCA1/2 Mutant Patients

To evaluate the relationship between gene expression patterns and FGA in *BRCA1/BRCA2*-mutant breast tumors, publicly available datasets from *TCGA PanCancer Atlas* and *TCGA Nature 2012* were accessed *via* cBioPortal, as specified in the corresponding figure legends. *BRCA1/2* mutation status and FGA values were obtained as described above. Analyses were restricted to samples harboring *BRCA1* or *BRCA2* mutations; non-mutant cases were excluded by filtering the dataset to retain only mutation-positive samples. In the TCGA *PanCancer Atlas* cohort, this included a total of 24 *BRCA1-*mutant and 28 *BRCA2-*mutant samples while in the *TCGA Nature 2012* cohort, this yielded 15 *BRCA1*-mutant and 22 *BRCA2*-mutant samples. To obtain gene expression data for these selected cases, a user-defined case list was generated in cBioPortal by inputting the corresponding patient identifiers. The mRNA expression levels were retrieved as z-scores relative to diploid samples, derived from RNA sequencing (RNA-Seq V2 RSEM) for the *TCGA PanCancer Atlas* dataset or microarray platforms for the *TCGA Nature 2012* dataset. Expression data were visualized using the heatmap function within the “Tracks” module and downloaded for downstream analyses; patient-level gene expression z-scores were exported and matched to corresponding FGA values. Samples were subsequently stratified into high and low expression groups using a threshold of z-score > 0 (high expression) and z-score < 0 (low expression). Associations between gene expression status and FGA were visualized and analyzed using GraphPad Prism (version 10.6.1). Statistical methods and sample sizes are detailed in the respective figure legends.

### Stratification by Tumor Stage and Molecular Subtype Relative to FGA

To assess the relationship between tumor stage, molecular subtype, and FGA, data from the *TCGA PanCancer Atlas* breast invasive carcinoma cohort were obtained via cBioPortal. FGA values were extracted for all available samples irrespective of *BRCA1/2* mutation status, as described above. Gene expression data were retrieved by querying genes of interest across all samples. The mRNA expression z-scores relative to diploid samples (RNA-Seq V2 RSEM) were visualized using the heatmap function within the “Tracks” module. Clinical annotations, including tumor stage and molecular subtype, were simultaneously obtained by selecting “American Joint Committee on Cancer Tumor Stage Code” and “Subtype” from the “Clinical” data tracks. Downloaded datasets were processed on a per-gene basis. For each gene, samples with low expression (z-score < 0) were excluded, and subsequent analyses were restricted to cases with high expression (z-score > 0). Corresponding FGA values for these samples were then analyzed. For downstream stratification, samples were grouped according to tumor stage (T classification) and molecular subtype. Tumor stage classification followed the TNM system based on primary tumor size: T1 (≤ 2 cm; including T1, T1a, T1b, T1c), T2 (> 2 cm to ≤ 5 cm), T3 (> 5 cm), and T4 (tumors with direct extension to the chest wall and/or skin, including inflammatory breast cancer subtypes such as T4d). Subcategories within T2 and T3 were consolidated due to limited sample representation. Molecular subtype analyses excluded samples classified as “Normal-like.” FGA distributions were subsequently compared across tumor stage and subtype groups for each gene. Data visualization and statistical analyses were performed using GraphPad Prism (version 10.6.1), with specific tests and sample sizes detailed in the corresponding figure legends.

### Identification of Mutation Burden of Different DDR and RST-associated Genes

To characterize the mutation burden of genes involved in DDR and RST pathways, data from the *TCGA PanCancer Atlas* breast invasive carcinoma cohort were obtained via cBioPortal. All available samples were included, and genes of interest were queried using the “Query by Gene” function, as previously described. Somatic mutation data were retrieved by selecting “Mutation count” within the “Tracks” module. Patient-level mutation counts were downloaded for downstream analysis. Samples with a mutation count of zero were excluded to focus on mutation-positive cases. The overall mutation burden across the cohort (“All”) was defined as the distribution of mutation counts across all mutation-positive samples, irrespective of gene-specific mutation status, and was used as a reference. For each gene of interest, samples harboring mutations in that gene were identified as a subset of the overall cohort. Mutation counts for gene-specific subsets were extracted and log10-transformed prior to analysis. These values were compared to the overall mutation burden distribution to assess relative mutation enrichment. Data visualization and statistical analyses were performed using GraphPad Prism (version 10.6.1), with specific statistical tests described in the corresponding figure legends.

### Identification of Co-occurring and Mutually Exclusive DDR Genes with Top Mutated Genes in Breast Cancer

To determine their co-occurring or mutually exclusive partners among our list of DDR-associated genes of interest, we again selected the the TCGA *PanCancer Atlas* cohort and included each of the top 6 mutated genes (separately) along with genes involved in different DDR pathways-assoaciated genes and “queried by gene” as previously described. We then navigated to the “Mutual Exclusivity” tab before downloading the data and plotting Log2Odds Ratio for each gene (top mutated) and its partners (DDR-associated genes) on GraphPad Prism.To quantify the mutation frequency of highly recurrently altered genes in basal-like breast cancer, data from the *TCGA PanCancer Atlas* breast invasive carcinoma cohort were accessed via cBioPortal, by exploring the selected study. The dataset was filtered to include only basal-like tumors by selecting the “Subtype” filter and restricting analyses to samples classified as “Basal” (n = 171). Mutation data for the most frequently altered genes (*TP53*, *BRCA1*, *PIK3CA*, *BRCA2*, *FANCD2*, and *BAP1*) were obtained from the “Mutated Genes” module and exported for downstream analysis. The percentage of samples harboring mutations in each gene was calculated and visualized. To assess patterns of co-occurrence and mutual exclusivity between these top mutated genes and DDR-associated genes, pairwise analyses were conducted using the cBioPortal “Mutual Exclusivity” module. Each of the six highly mutated genes was queried individually in combination with a curated list of DDR pathway genes. Statistical associations between gene pairs were quantified using log2 odds ratios, as provided by cBioPortal. Downloaded data were processed and visualized using GraphPad Prism (version 10.6.1). Log2 odds ratios were plotted to represent the degree of co-occurrence (positive values) or mutual exclusivity (negative values) between gene pairs. Statistical significance and sample sizes are reported in the corresponding figure legends.

### Statistical Analysis

All stastistical tests were performed on GraphPad Prism (version 10.6.1). Comparisons between two groups (e.g., *BRCA1/2* mutant versus wild-type, or high versus low gene expression) were conducted using a two-tailed unpaired Student’s *t*-test. For analyses involving more than two groups (e.g., tumor stage, molecular subtype, or gene-specific mutation burden relative to the overall cohort), differences in mean Fraction of Genome Altered (FGA) were evaluated using one-way analysis of variance (ANOVA), followed by Tukey’s post hoc test for multiple pairwise comparisons. Co-occurrence and mutual exclusivity of genomic alterations were assessed using data obtained from cBioPortal where statistical significance was determined using two-sided Fisher’s exact test. All statistical tests were two-sided, and a *P* value < 0.05 was considered statistically significant. Specific statistical tests, sample sizes, and exact *P* values are provided in the corresponding figure legends.

## Data Availability

The data in this study are available within the article and upon request.

## Acknowledgements

We thanks to Dr. Lixing Yang and Dr. Yang Yang for helpful discussions. This work was supported in part by Mayo Clinic Breast Cancer SPORE grant P50 CA116201 from National Institute of Health (to A.P.), R00CA252044-03 (to A.P.)

## Author Contributions

**Conceptualization:** Farah Ramadan and Arvind Panday

**Data Curation:** Farah Ramadan, Sara Zahraeifard, Ujwal Subedi, Neelima Yadav, and Arvind Panday

**Methodology**: Sara Zahraeifard, Farah Ramadan, Neelima Yadav

**Formal Analysis:** Farah Ramadan and Arvind Panday

**Funding Acquisition**: Arvind Panday

**Supervision:** Arvind Panday

**Writing – original draft**: Arvind Panday and Farah Ramadan.

**Writing – review & editing**: Farah Ramadan, Sara Zahraeifard, Sameer Shah and Arvind Panday

## Materials & Correspondence

All correspondence should be addressed to A.P.

## SUPPLEMENTARY FIGURE LEGENDS

**Supplementary Figure S1. BRCA2 deficiency drives genomic instability and adaptive DNA repair pathway activation in breast tumors (A)** FGA relative to *BRCA2* gene status (Wild-type/Mutant) from three different Invasive Breast Carcinoma studies including *MSK 2025* (n = 3841) **(left)**, TCGA *Nature 2012* (n = 482) **(middle)** and TCGA *PanCancer Atlas* (n = 1050) **(right)**. **(B-G)** FGA relative to expression levels of **(B)** HR genes in *BRCA2*-mutant samples; **(C)** NHEJ genes in *BRCA1*-mutant **(upper panel)** or *BRCA2*-mutant **(lower panel)** samples; **(D)** FA genes, **(E)** MMR genes, **(F)** aEJ genes, and **(G)** BER genes, all in *BRCA2-*mutant breast cancer samples. *BRCA1*-mutant (n = 24) and *BRCA2*-mutant (n = 28) patients are coming from TCGA *PanCancer Atlas* study (except XRCC1 coming from TCGA *Nature 2012* (n= 22)). Low expression (heatmap mRNA expression z-score <0) and High expression (heatmap mRNA expression z-score >0). Asterisks denote significant p-value using two-tailed unpaired t-test with ***p<0.001, **p<0.01, *p<0.05; ns = non-significant.

**Supplementary Figure S2. Replication stress tolerance pathways are selectively upregulated in *BRCA2* mutant breast tumors**. FGA relative to expression level of genes involved in different stress tolerance mechanisms including **(A)** Fork remodeling/reversal, **(B)** Fork protection, **(C)** ssDNA gap suppression**, (D)** Replication Fidelity and **(E)** Mitotic DNA synthesis, in *BRCA2-*mutant breast cancer patients. Low expression (heatmap mRNA expression z-score < 0) and High expression (heatmap mRNA expression z-score > 0). All patients shown belong to the TCGA *PanCancer Atlas* cohort (n = 28). Asterisks denote significant p-value using two-tailed unpaired t-test with ***p<0.001, **p<0.01, *p<0.05; ns = non-significant. RAD52 is shown in two different studies: TCGA *PanCancer Atlas* (ns) and TCGA *Nature 2012* where indicated (n = 22).

