## Supplementary figures and images for "Comprehensive Analysis Reveals Adaptive DNA Repair and Replication Stress Networks in Genomically Unstable Breast Cancer"

### Supplement data

#### Supplementary Figure S1

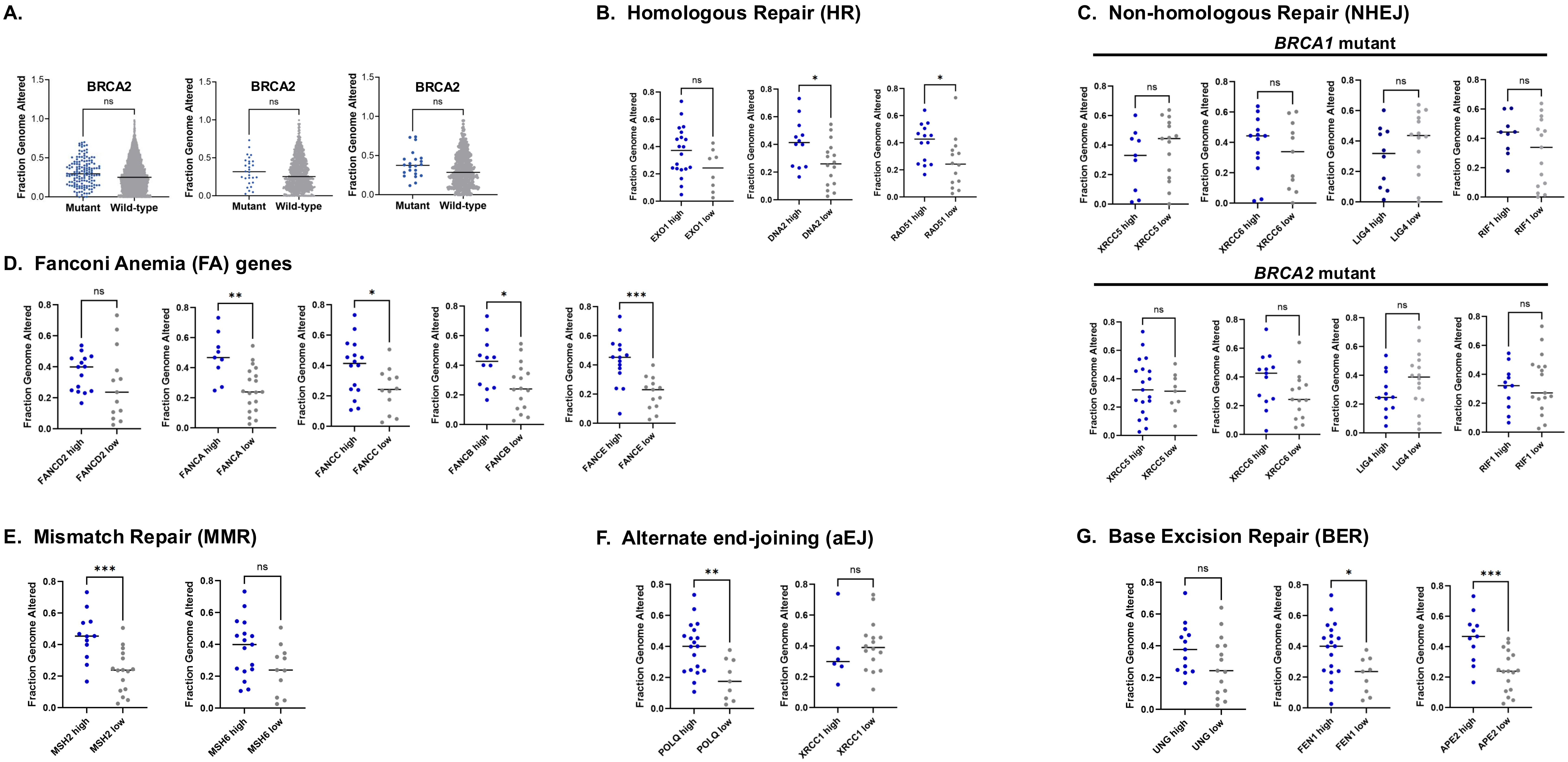

#### Supplementary Figure S2

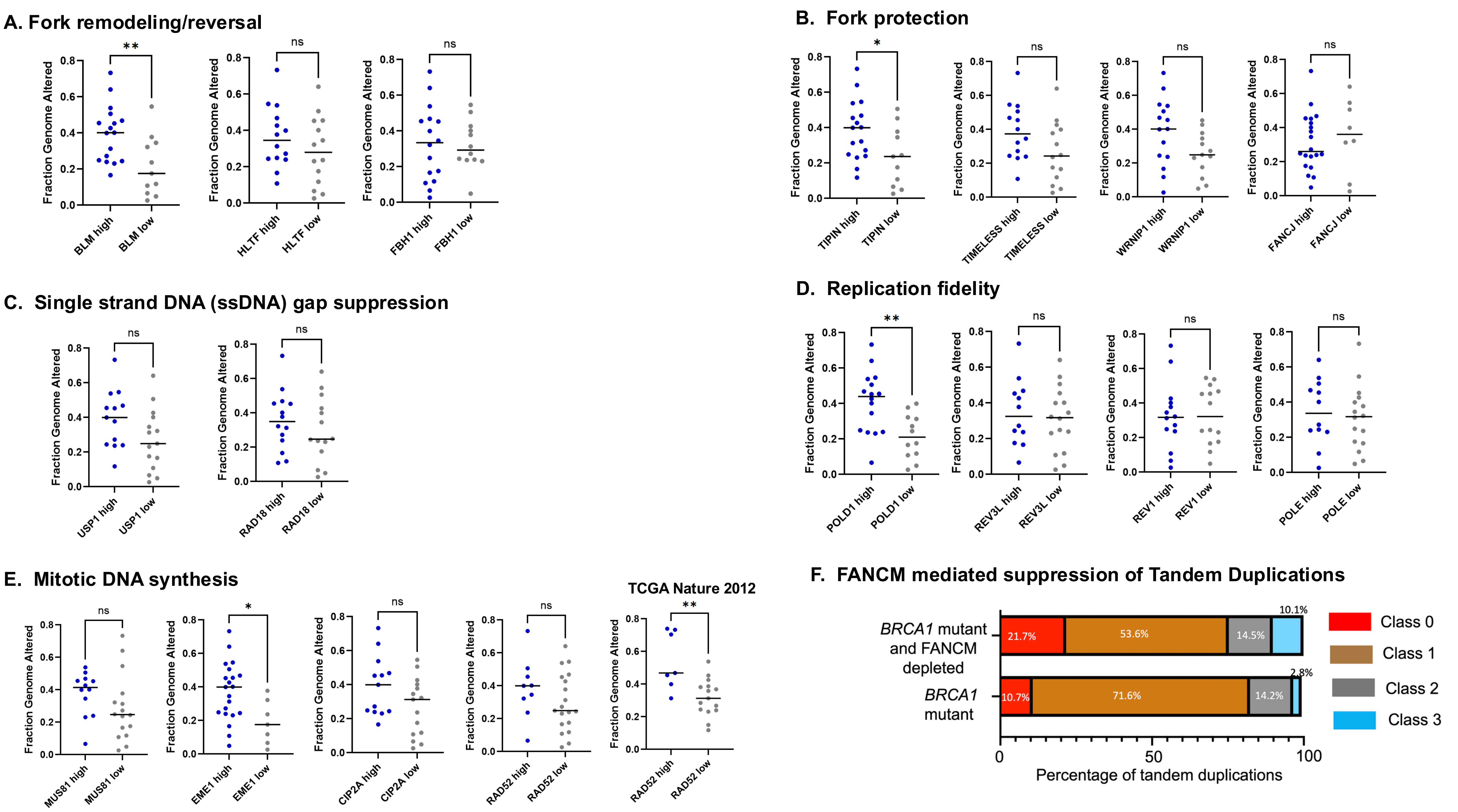
